# Viral coinfection is shaped by host ecology and virus-virus interactions across diverse microbial taxa and environments

**DOI:** 10.1101/038877

**Authors:** Samuel L. Díaz Muñoz

**Affiliations:** Center for Genomics and Systems Biology + Department of Biology 12 Waverly Place New York University, New York, NY, USA 10003

## Abstract

Infection of more than one virus in a host, coinfection, is common across taxa and environments. Viral coinfection can enable genetic exchange, alter the dynamics of infections, and change the course of viral evolution. Yet, a systematic test of the factors explaining variation in viral coinfection across different taxa and environments awaits completion. Here I employ three microbial data sets of virus-host interactions covering cross-infectivity, culture coinfection, and single-cell coinfection (total: 6,564 microbial hosts, 13,103 viruses) to provide a broad, comprehensive picture of the ecological and biological factors shaping viral coinfection. I found evidence that ecology and virus-virus interactions are recurrent factors shaping coinfection patterns. Host ecology was a consistent and strong predictor of coinfection across all three datasets: cross-infectivity, culture coinfection, and single-cell coinfection. Host phylogeny or taxonomy was a less consistent predictor, being weak or absent in the cross-infectivity and single-cell coinfection models, yet it was the strongest predictor in the culture coinfection model. Virus-virus interactions strongly affected coinfection. In the largest test of superinfection exclusion to date, prophage sequences reduced culture coinfection by other prophages, with a weaker effect on extrachromosomal virus coinfection. At the single-cell level, prophage sequences eliminated coinfection. Virus-virus interactions also *increased* culture coinfection with ssDNA-dsDNA coinfections >2x more likely than ssDNA-only coinfections. The presence of CRISPR spacers was associated with a ~50% reduction in single-cell coinfection in a marine bacteria, despite the absence of exact spacer matches in any active infection. Collectively, these results suggest the environment bacteria inhabit and the interactions among surrounding viruses are two factors consistently shaping viral coinfection patterns. These findings highlight the role of virus-virus interactions in coinfection with implications for phage therapy, microbiome dynamics, and viral infection treatments.

## Introduction

Viruses outnumber hosts by a significant margin (Bertani, 1954; Bergh *et al.,* 1989; Suttle, 2007; Weinbauer, 2004; Rohwer and Barott, 2012; Wigington *et al.,* 2016). In this situation, infection of more than one strain or type of virus in a host (coinfection) might be expected to be a rather frequent occurrence potentially leading to virus-virus interactions (Berngruber *et al.,* 2010; Díaz-Muñoz and Koskella, 2014; Bergh *et al.,* 1989; Rohwer and Barott, 2012; Suttle, 2007; Weinbauer, 2004). Across many different viral groups, virus-virus interactions within a host can alter genetic exchange (Dang *et al.,* 2004; Worobey and Holmes, 1999; Cicin-Sain *et al.,* 2005; Turner *et al.,* 1999), modify viral evolution (Joseph *et al.,* 2009; Refardt, 2011; Roux *et al.,* 2015; Dropulić *et al.,* 1996; Ghedin *et al.,* 2005; Turner and Chao, 1998), and change the fate of the host (Roux *et al.,* 2012; Vignuzzi *et al.,* 2006; Li *et al.,* 2010; Abrahams *et al.,* 2009). Yet, there is little information regarding the ecological and biological factors that shape coinfection and virus-virus interactions. Given that most laboratory studies of viruses focus on a single virus at a time (DaPalma *et al.,* 2010), understanding the drivers of coinfection and virus-virus interactions is a pressing frontier for viral ecology.

Recent studies of bacteriophages have highlighted just how widespread coinfection is and have started to shed light on the ecology of viral coinfection. For example, although studies of phage host range have shown that some bacterial hosts can be infected by more than one type of phage for some time (Moebus and Nattkemper, 1981; Kaiser and Dworkin, 1975; Spencer, 1957), mounting evidence indicates this is a pervasive phenomenon (Koskella and Meaden, 2013; Flores *et al.,* 2013; 2011). The ability of two or more viruses to independently infect the same host –cross-infectivity– revealed in these studies of host range is a necessary, but not sufficient criterion to determine coinfection. Thus, cross-infectivity or the potential for coinfection, represents a baseline shaping coinfection patterns. To confirm coinfection, sequence-based approaches have been used to detect viral coinfection at two scales: >1 virus infecting the same multicellular host (e.g., a multicellular eukaryote or a colony of bacterial cells, hereafter called culture coinfection), or to the coinfection of a single cell. At the culture coinfection scale, a study mining sequence data to examine virus-host relationships uncovered widespread coinfection in publicly available bacterial and archaeal genome sequence data (Roux *et al.,* 2015). At the single-cell scale, two studies have provided, for the first time, single-cell level information on viruses associated with specific hosts isolated from the environment in a culture-independent manner (Roux *et al.,* 2014; Labonté *et al.,* 2015). Collectively, these studies suggest that there is a large potential for coinfection and that this potential is realized at both the host culture and single cell scales. A summary of these studies suggests roughly half of hosts have the potential to be infected (i.e. are cross-infective) or are actually infected by an average of >2 viruses (Table 1). Thus, there is extensive evidence across various methodologies, taxa, environments, that coinfection is widespread and virus-virus interactions may be a frequent occurrence.

**Table 1.** Viral coinfection is prevalent across various methodologies, taxa, environments, and scales of coinfection.

|  | Cross-infectivity<br>(*potential* coinfection) | Culture<br>coinfection | Single-cell<br>coinfection |
| --- | --- | --- | --- |
| Number of viruses | 4.89 (± 4.61) | 3.377 ± 1.804 | 2.37 ± 0.83 |
| Prop of bacteria w/ multiple infections | 0.654 | 0.538 | 0.450 |
| Reference | (GOLD: Mukherjee *et al.*, 2017; Flores *et al.*, 2011) | (Roux *et al.*, 2014; 2015) | (Roux *et al.*, 2014) |

The aforementioned studies, among others, establish that viral coinfection is a frequent occurrence in bacterial and archaeal hosts, but systematic tests of the factors explaining variation in viral coinfection across different taxa and environments are lacking. However, the literature suggests four factors that are likely to play a role: host ecology, host taxonomy or phylogeny, host defense mechanisms, and virus-virus interactions. The relevance and importance of these are likely to vary for cross-infectivity, culture coinfection, and single-cell coinfection.

Cross-infectivity has been examined in studies of phage host-range, which have provided some insight into the factors affecting potential coinfection. In a single bacterial species there can be wide variation in phage host range (Holmfeldt *et al.,* 2007), and thus, cross-infectivity. A larger, quantitative study of phage-bacteria infection networks in multiple taxa also found wide variation in cross-infectivity in narrow taxonomic ranges (strain or species level), with some hosts susceptible to few viruses and others to many (R Development Core Team, 2011). However, at broader host taxonomic scales, cross-infectivity followed a modular pattern, suggesting that higher taxonomic ranks could influence cross-infectivity (Flores *et al.,* 2013). Thus, host taxonomy may show a scale dependent effect in shaping coinfection patterns. Flores et al. (2013) also highlighted the potential role of ecology in shaping coinfection patterns as geographic separation played a role in cross-infectivity. Thus, bacterial ecology and phylogeny (particularly at taxonomic ranks above species) are the primary candidates for drivers of coinfection.

Culture and single cell coinfection, require cross-infectivity, but also require both the bacteria and infecting viruses to allow simultaneous or sequential infection. Thus, in addition to bacterial phylogeny and ecology, we can expect additional factors shaping coinfection, namely bacterial defense mechanisms and virus-virus interactions. An extensive collection of studies provides evidence that bacterial and viral mechanisms may affect coinfection. Bacteria, understandably reluctant to welcome viruses, possess a collection of mechanisms of defense against viral infection (Labrie *et al.,* 2010), including restriction enzymes (Murray, 2002; Linn and Arber, 1968) and CRISPR-Cas systems (Horvath and Barrangou, 2010). The latter have been shown to be an adaptive immune system for bacteria, protecting from future infection by the same phage (Barrangou *et al.,* 2007) and preserving the memory of viral infections past (Held and Whitaker, 2009). Metagenomic studies of CRISPR in natural environments suggest rapid coevolution of CRISPR arrays (Tyson and Banfield, 2008), but little is known regarding the *in-situ* protective effects of CRISPR on cells, which should now be possible to decipher with single-cell genomics.

Viruses also have mechanisms to mediate infection by other viruses, some of which were identified in early lab studies of bacteriophages (Ellis and Delbruck, 1939; Delbruck, 1946). An example of a well-described phenomenon of virus-virus interactions is superinfection immunity conferred by lysogenic viruses (Bertani, 1953), which can inhibit coinfection of cultures and single cells (Bertani, 1954). While this mechanism has been described in several species, its frequency at broader taxonomic scales and its occurrence in natural settings is not well known.

Most attention in virus-virus interactions has focused on mechanisms limiting coinfection, with the assumption that coinfection invariably reduces host fitness (Berngruber *et al.,* 2010). However, some patterns of non-random coinfection suggest elevated coinfection (Dang *et al.,* 2004; Cicin-Sain *et al.,* 2005; Turner *et al.,* 1999) and specific viral mechanisms that promote co-infection have been identified, including enhanced expression of the phage attachment site upon infection (Joseph *et al.,* 2009), particle aggregation (Altan-Bonnet and Chen, 2015; Aguilera *et al.,* 2017), and phage-phage communication (Erez *et al.,* 2017). Systematic coinfection has been proposed (Roux *et al.,* 2012) to explain findings of chimeric viruses of mixed nucleic acids in metagenome reads (Diemer and Stedman, 2012; Roux *et al.,* 2013). This suspicion was confirmed in a study of marine bacteria that found highly non-random patterns of coinfection between ssDNA and dsDNA viruses in a lineage of marine bacteria (Roux *et al.,* 2014), but the frequency of this phenomenon across bacterial taxa remains to be uncovered. Thus, detailed molecular studies of coinfection dynamics and virome sequence data are generating questions ripe for testing across diverse taxa and environments.

Here I employ virus-host interaction data sets to date to provide a broad, comprehensive picture of the ecological and biological factors shaping coinfection. The data sets are each the largest of their kind, providing an opportunity to examine viral coinfection at multiple scales, from cross-infectivity (1,005 hosts; 499 viruses), to culture coinfection (5,492 hosts; 12,498 viruses) and single-cell coinfection (127 hosts; 143 viruses) to answer the following questions:

1. How do ecology, bacterial phylogeny/taxonomy, bacterial defense mechanisms, and virus-virus interactions explain variation in estimates of viral coinfection?
2. What is the relative importance of each of these factors in cross-infectivity, culture coinfection, and single-cell coinfection?
3. Do prophage sequences limit viral coinfection?
4. Do ssDNA and dsDNA viruses show evidence of preferential coinfection?
5. Do sequences associated with the CRISPR-Cas bacterial defense mechanism limit coinfection of single cells?

The results of this study suggest that microbial host ecology and virus-virus interactions are consistently important mediators of the frequency and extent of coinfection. Host taxonomy and CRISPR spacers also shaped culture and single-cell coinfection patterns, respectively. Virus-virus interactions served to limit and promote coinfection.

## Materials and Methods

### Data Sets

I assembled data collectively representing 13,103 viral infections in 6,564 bacterial and archaeal hosts from diverse environments (Table S1). These data are composed of three data sets that provide an increasingly fine-grained examination of coinfection from cross-infectivity (potential coinfection), to coinfection at the culture (pure cultures or single colonies, not necessarily single cells) and single-cell levels. The data set examining cross-infectivity assessed infection experimentally with laboratory cultures, while other two data sets (culture and single-cell coinfection) used sequence data to infer infection.

The first data set on cross-infectivity (potential coinfection) is composed of bacteriophage host-range infection matrices documenting the results of experimental attempts at lytic infection in cultured phage and hosts (Flores *et al.,* 2011). It compiles results from 38 published studies, encompassing 499 phages and 1,005 bacterial hosts. Additionally, I entered new metadata (ecosystem and ecosystem category) to match the culture coinfection data set (see description below), to enable comparisons between these two multi-taxon data sets. The host-range infection data are matrices of infection success or failure via the “spot test”, briefly, a drop of phage lysate is “spotted” on a bacterial lawn and lysing of bacteria is noted as presence or absence. This data set represents studies with varying sample compositions, in terms of bacteria and phage species, bacterial energy sources, source of samples, bacterial association, and isolation habitat.

The second data set on culture coinfection is derived from viral sequence information mined from published microbial genome sequence data on National Center for Biotechnology Information’s (NCBI) databases (Roux *et al.,* 2015). Thus, this second data set provided information on actual (as opposed to potential) coinfection of cultures, representing 12,498 viral infections in 5,492 bacterial and archaeal hosts. The set includes data on viruses that are incorporated into the host genome (prophages) as well as extrachromosomal viruses detected in the genome assemblies (representing lytic, chronic, carrier state, and ‘extrachromosomal prophage’ infections). Genomes of microbial strains were primarily generated from colonies or pure cultures (except for 27 hosts known to be represented by single cells). Thus, although these data could represent coinfection at the single-cell level, they are more conservatively regarded as culture coinfections. In addition, I imported metadata from the US Department of Energy, Joint Genome Institute, Genomes Online Database (GOLD: Mukherjee *et al.,* 2017) to assess the ecological and host factors (ecosystem, ecosystem category, and energy source) that could influence culture coinfection. I curated these records and added missing metadata (n = 4964 records) by consulting strain databases (e.g. NCBI BioSample, DSMZ, BEI Resources, PATRIC and ATCC) and the primary literature.

The third data set included single-cell amplified genomes, providing information on coinfection and virus-virus interactions within single cells (Roux *et al.,* 2014). This genomic data set covers viral infections in 127 single cells of SUP05 marine bacteria (sulfur-oxidizing Gammaproteobacteria) isolated from different depths of the oxygen minimum zone in the Saanich Inlet near Vancouver Island, British Columbia (Roux *et al.,* 2014). These single-cell data represent a combined 143 viral infections including past infections (CRISPRs and prophages) and active infections (active at the time of isolation, e.g. ongoing lytic infections).

A list and description of data sources are included in Supplementary Table 1, and the raw data used in this paper are deposited in the FigShare data repository (FigShare doi:10.6084/m9.figshare.2072929).

### Factors explaining cross-infectivity and coinfection

To test the factors that potentially influence coinfection –ecology, host taxonomy/phylogeny, host defense mechanisms, and virus-virus interactions– I conducted regression analyses on each of the three data sets, representing an increasingly fine scale analysis of coinfection from cross-infectivity, to culture coinfection, and finally to single-cell coinfection. The data sets represent three distinct phenomena related to coinfection and thus the variables and data in each one are necessarily different. The measures of coinfection, for example, were 1) the amount of phage that could infect each host, 2) the number of extrachromosomal infections, and 3) the number of active viral infections, respectively, for cross-infectivity, culture coinfection, and single-cell coinfection data sets. Virus-virus interactions were measured as the association of these aforementioned ongoing viral infections with the presence of other types of viruses (e.g. prophages) or the association of viruses with different characteristics (e.g. ssDNA vs dsDNA). Ecological factors and bacterial taxonomy were tested in all data sets, but virus-virus interactions, for example, were not evaluated in the cross-infectivity data set because they do not apply (i.e., measures potential and not actual coinfection). Table 2 details the data used to test each of the factors that potentially influence coinfection and details on the analysis of each data set are provided in turn below.

**Table 2.**
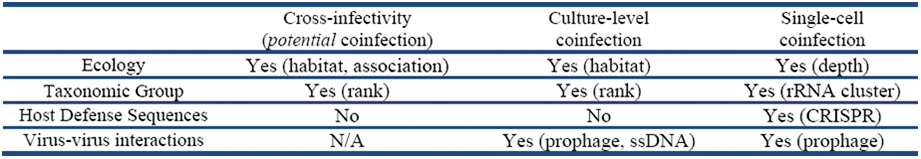
Factors explaining coinfection tested with each of the three data sets and (specific variable tested).

First, to test the potential influence of these factors on cross-infectivity, I conducted a factorial analysis of variance (ANOVA) on the cross-infectivity data set. The dependent variable was the amount of phage that could infect each host, measured specifically as the proportion of tested phage that could infect each host because the infection matrices in this data set were derived from different studies testing varying numbers of phages. Thus, as the number of phage that could infect each host (cross-infectivity) increased, the potential for coinfection was greater. The five independent variables tested were the study type/source (natural, coevolution, artificial), bacterial taxon (roughly corresponding to Genus), habitat from which bacteria and phages were isolated, bacterial energy source (photosynthetic or heterotrophic), and bacterial association (e.g. pathogen, free-living). Geographic origin and phage taxa were present in original the metadata, but were largely incomplete; therefore they were not included in the analyses. Model criticism suggested that ANOVA assumptions were reasonably met (Supplementary Figure 1), despite using proportion data (Warton and Hui, 2011). ANOVA on the arc-sine transform of the proportions and a binomial regression provided qualitatively similar results (data not shown, see associated code in FigShare repository).

**Figure 1.**
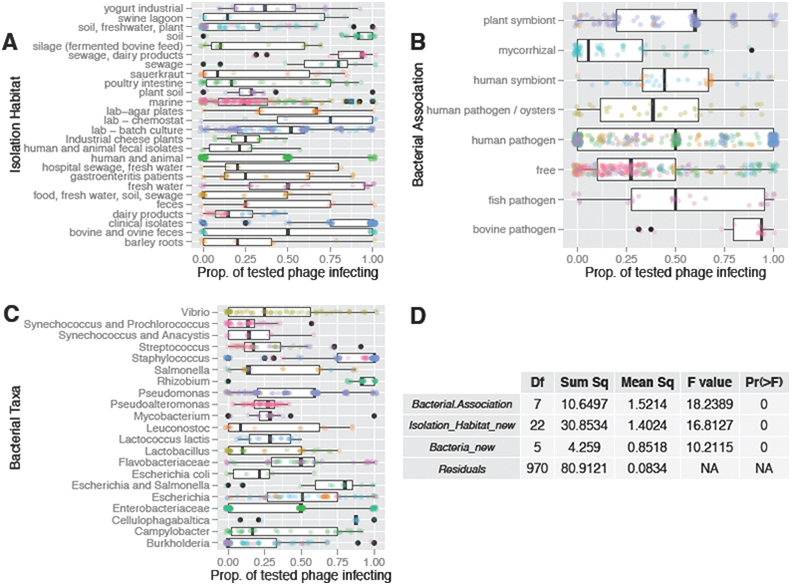
Bacterial-phage ecology and taxonomy explain most of the variation in cross-infectivity. The data set is a compilation of 38 phage host range studies that measured the ability of different phages (n=499) to infect different bacterial hosts (n=1,005) using experimental infections of cultured microbes in the laboratory (i.e., the “spot test”). Cross-infectivity, or potential coinfection, is the number of phages that can infect a given bacterial host, here measured as the proportion of tested phages infecting each host (represented by points). Cross-infectivity was the response variable in an analysis of variance (ANOVA) that examined the effect of five factors (study type, bacterial taxon, habitat from which bacteria and phages were isolated, bacterial energy source, and bacterial association) on variation in the response variable. Those factors selected after stepwise model selection using Akaike’s Information Criterion (AIC) are depicted in panels A-C. In these panels, each point represents a bacterial host; hosts with a value of zero on the x-axis could be infected by none of the tested phage, whereas those with a value of one could be infected by all the tested phages. Thus, the x-axis is a scale of the potential for coinfection. Point colors correspond to hosts that were tested in the same study. Note data points are offset by a random and small amount (jittered) to enhance visibility and reduce overplotting. The ANOVA table is presented in panel D.

Second, to test the factors influencing culture coinfection I conducted a negative binomial regression on the culture coinfection data set. The number of extrachromosomal viruses infecting each host culture, as detected by sequence data, was the dependent variable. These viruses represent ongoing infections (e.g. lytic, chronic, or carrier state), as opposed to viral sequences integrated into the genome that may or may not be active. Thus, as more extrachromosomal infections are detected in a host culture, culture coinfection increases. The five explanatory variables tested were the number of prophage sequences, ssDNA virus presence, energy source (e.g. heterotrophic/autotrophic), taxonomic rank (Genus was selected as the best predictor over Phylum or Family, details in code in Figshare repository), and habitat (environmental, host-associated, or engineered; as defined by GOLD specifications (Mukherjee *et al.,* 2017)). I conducted stepwise model selection using Akaike’s Information Criterion (AIC), a routinely used measure of entropy that ranks the relative quality of alternative models (Akaike, 1974), to arrive at a reduced model that minimized the deviance of the regression.

Third, to test the factors influencing single cell coinfection I conducted a Poisson regression on single cell data set. The number of actively infecting viruses in each cell, as detected by sequence data, was the dependent variable. This variable excludes viral sequences integrated into the genome (e.g. prophages) that may or may not be active. Thus, as the number of active viral infections detected increase, single cell coinfection is greater. The four the explanatory variables tested were phylogenetic cluster (based on small subunit rRNA amplicon sequencing), ocean depth, number of prophage sequences, and number of CRISPR spacers. I conducted stepwise model selection with AIC (as done with the culture coinfection data set, see above) to arrive at a reduced model that minimized deviance.

### Virus-virus interactions (prophages and ss/dsDNA) in culture and single cell coinfection

To obtain more detailed information (beyond the regression analyses) on the influence of virus-virus interactions on the frequency and extent of coinfection, I analyzed the effect of prophage sequences in culture and single-cell coinfections, and the influence of ssDNA and dsDNA viruses on culture coinfection.

The examinations of prophage infections were entirely sequence-based in the culture coinfection and single-cell coinfection data sets. Because only sequence data is used, the activity of these prophage sequences cannot be confirmed. For instance, the prophage sequences in the original single-cell data set were conservatively termed ‘putative defective prophages’ (Roux *et al.,* 2014), because sequences were too small to be confidently assigned as complete prophage sequences (S. Roux, pers. comm.). Therefore, I will hereafter be primarily using the phrase ‘prophage sequences’ (especially when indicating results) synonymously with ‘prophages’, primarily for grammatical convenience. To determine whether prophage sequences affected the frequency and extent of coinfection in the culture coinfection data set, I examined host cultures infected exclusively by prophage sequences or extrachromosomal viruses (representing chronic, carrier state, and ‘extrachromosomal prophage’ infections) and all prophage-infected cultures. I tested whether prophage-only coinfections were infected by a different average number of viruses compared to extrachromosomal-only coinfections using a Wilcoxon Rank Sum test. I tested whether prophage-infected cultures were more likely than not to be coinfections, and whether these coinfections were more likely to occur with additional prophage sequences or extrachromosomal viruses.

To examine the effect of prophage sequences on coinfection frequency in the single-cell data set, I examined cells with prophage sequences and calculated the proportion of those that also had active infections. To determine the extent of coinfection in cells with prophage sequences, I calculated the average number of active viruses infecting these cells. I examined differences in the frequency of active infection between bacteria with or without prophage sequences using a proportion test and differences in the average amount of current viral infections using a Wilcoxon rank sum test.

To examine whether ssDNA and dsDNA viruses exhibited non-random patterns of culture coinfection, as found by Roux et al. (2014) for single cells of SUP05 marine bacteria, I examined the larger culture coinfection data set (based on Roux *et al.,* 2015) that is composed of sequence-based viral detections in all bacterial and archaeal NCBI genome sequences. I first compared the frequency of dsDNA-ssDNA mixed coinfections against ssDNA-only coinfections among all host cultures coinfected with at least one ssDNA virus (n=331), using a binomial test. To provide context for this finding (because ssDNA viruses were a minority of detected viral infections) I also examined the prevalence of dsDNA-only and dsDNA-mixed coinfections and compared the proportion of ssDNA-mixed/total-ssDNA coinfections with the proportion of dsDNA-mixed/dsDNA-total coinfections using a proportion test. To examine whether potential biases in the detectability of ssDNA viruses relative to dsDNA viruses (Roux *et al.,* 2016) could affect ssDNA-dsDNA coinfection detection, I also tested the percent detectability of ssDNA viruses in the overall data set, compared to the percentage of coinfected host cultures that contained at least one ssDNA virus using a proportion test. Finally, because the extended persistent infections maintained by some ssDNA viruses (particularly in the *Inoviridae* family) have been proposed to underlie previous patterns of enhanced coinfection between dsDNA and ssDNA viruses, I examined the proportion of coinfections involving ssDNA viruses that had at least one virus of the *Inoviridae* among ssDNA-only coinfections, ssDNA single infections, and ssDNA-dsDNA coinfections.

### Effect of CRISPR spacers on single cell coinfection

In order to obtain a more detailed picture of the potential role of bacterial defense mechanisms in coinfection, I investigated the effect of CRISPR spacers (sequences retained from past viral infections by the CRISPR-Cas defense mechanisms) on the frequency of cells undergoing active infections in the single cell data set (n=127 cells of SUP05 bacteria). I examined cells that harbored CRISPR spacers in the genome and calculated the proportion of those that also had active infections. To determine the extent of coinfection in cells with CRISPR spacers, I calculated the average number of active viruses infecting these cells. I examined differences in the frequency of cells undergoing active infections in cells with or without CRISPR spacers using a proportion test. I examined differences in the average number of current viral infections in cells with or without CRISPR spacers using a Wilcoxon rank sum test.

### Statistical Analyses

I conducted all statistical analyses in the R statistical programming environment (R Development Core Team, 2011) and generated graphs using the ggplot2 package (Wickham, 2009). Means are presented as means ± standard deviation, unless otherwise stated. Data and code for analyses and figures are available in the Figshare data repository (FigShare doi:10.6084/m9.figshare.2072929).

## Results

### Host ecology and virus-virus interactions consistently explain variation in potential (cross-infectivity), culture, and single-cell coinfection

To test the factors that influence viral coinfection of microbes, I conducted regression analyses on each of the three data sets (cross-infectivity, culture coinfection, and single cell coinfection). The explanatory variables tested in each data set varied slightly (Table 2), but can be grouped into host ecology, virus-virus interactions, host taxonomic or phylogenetic group, and sequences associated with host defense. Overall, host ecology and virus-virus interactions (association of ongoing infections with other viruses, e.g. prophages or ssDNA viruses) appeared as statistically significant factors that explained a substantial amount of variation in coinfection in every data set they were tested (see details for each data set in subheadings below). Host taxonomy was a less consistent predictor across the data sets. The taxonomic group of the host explained little variation or was not a statistically significant predictor of cross-infectivity and single-cell coinfection, respectively. However, host taxonomy was the strongest predictor of the amount of culture coinfection, judging by the amount of deviance explained in the negative binomial regression. Sequences associated with host defense mechanisms, specifically CRISPR spacers, were tested only in the single cell data set (n = 127 cells of SUP05 marine bacteria) and were a statistically significant and strong predictor of coinfection.

### Potential for coinfection (cross-infectivity) is shaped by bacterial ecology and taxonomy

To test the viral and host factors that affected cross-infectivity, I conducted an analysis of variance on the cross-infectivity data set, a compilation of 38 phage host-range studies that measured the ability of different phages (n=499) to infect different bacterial hosts (n=1,005) using experimental infections of cultured microbes in the laboratory (Flores *et al.,* 2011). In this data set cross-infectivity is the proportion of tested phage that could infect each host. Stepwise model selection with AIC, yielded a reduced model that explained 33.89% of the variance in cross-infectivity using three factors: host association (e.g. free-living, pathogen), isolation habitat (e.g. soil, animal), and taxonomic grouping (Figure 1). The two ecological factors together explained >30% of the variance in cross-infectivity, while taxonomy explained only ~3% (Figure 1D). Host association explained 8.41% of the variance in cross-infectivity. Bacteria that were pathogenic to cows had the highest potential coinfection with more than 75% of tested phage infecting each bacterial strain (Figure 1B). In absolute terms, the average host that was a pathogenic to cows could be infected 15 phages on average. The isolation habitat explained 24.36% of the variation in cross-infectivity with clinical isolates having the highest median cross-infectivity, followed by sewage/dairy products, soil, sewage, and laboratory chemostats. All these habitats had more than 75% of tested phage infecting each host on average (Figure 1A) and, in absolute terms, the average host in each of these habitats could be infected by 3-15 different phages.

Bacterial energy source (heterotrophic, autotrophic) and the type of study (natural, coevolution, artificial) were not selected by AIC in final model. An alternative categorization of habitat that matched the culture coinfection data set (ecosystem: host-associated, engineered, environmental) was not a statistically significant predictor and was dropped from the model. Results from this alternative categorization (Figure S1 and Table S2) show that bacteria-phage groups isolated from host-associated ecosystems (e.g. plants, humans) had the highest cross-infectivity, followed by engineered ecosystems (18% lower), and environmental ecosystems (44% lower). Details of the full, reduced, and alternative models are provided in Figure S2 and associated code in the FigShare repository.

### Culture coinfection is influenced by host ecology and taxonomy and virus-virus interactions

To test the viral and host factors that affected culture coinfection, I conducted a negative binomial regression on the culture coinfection data set, which documented 12,498 viral infections in 5,492 microbial hosts using NCBI-deposited sequenced genomes (Roux *et al.,* 2015) supplemented with metadata collected from the GOLD database (Mukherjee *et al.,* 2017). Culture coinfection was measured as the number of extrachromosomal virus sequences (representing lytic, chronic, or carrier state infections) detected in each host culture. Stepwise model selection with AIC resulted in a reduced model that used three factors to explain the number of extrachromosomal infections: host taxonomy (Genus), number of prophages and, host ecosystem. The genera with the most coinfection (mean >2.5 extrachromosomal infections) were *Avibacterium, Shigella, Selenomonas, Faecalibacterium, Myxococcus,* and *Oenococcus,* whereas *Enterococcus, Serratia, Helicobacter, Pantoea, Enterobacter, Pectobacterium, Shewanella, Edwardsiella,* and *Dickeya* were rarely (mean < 0.45 extrachromosomal infections) coinfected (Figure 2A). Microbes that were host-associated had the highest mean extrachromosomal infections (1.39 ± 1.62, n=4,282), followed by engineered (1.32 ± 1.44, n=281) and environmental (0.95 ± 0.97, n=450) ecosystems (Figure 2B). The model predicts that each additional prophage sequence yields ~79% the amount of extrachromosomal infections, or a 21% reduction (Figure 2C).

Host energy source (heterotrophic, autotrophic) and ssDNA virus presence were not statistically significant predictors as judged by AIC model selection. Details of the full, reduced, and alternative models are provided in the Supplementary Materials and all associated code and data is deposited in FigShare repository.

**Figure 2.**
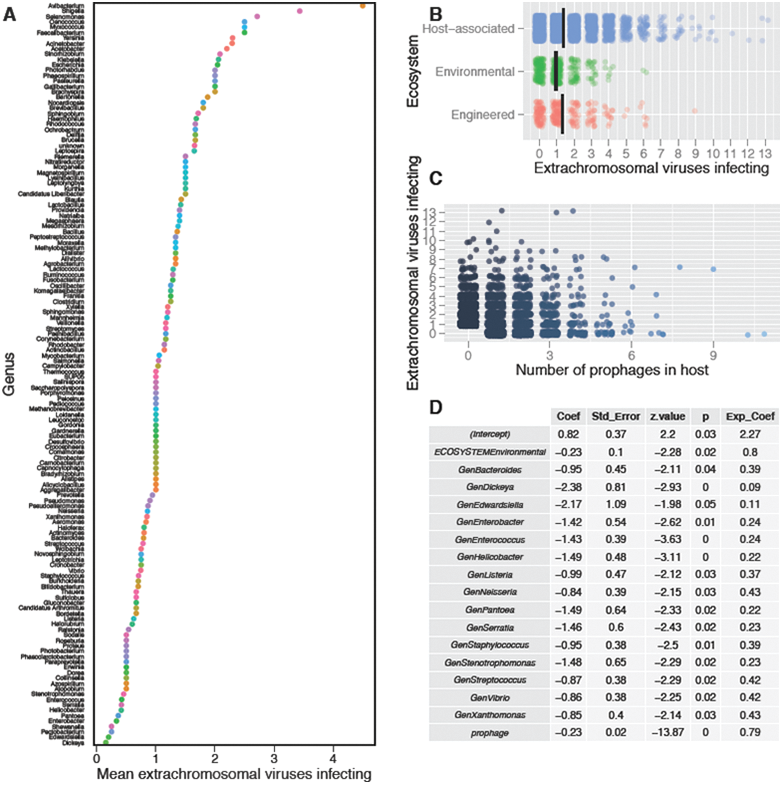
Host taxonomy, ecology, and number of prophage sequences predict variation in culture coinfection. The data set is composed of viral infections detected using sequence data (n=12,498) in all bacterial and archaeal hosts (n=5,492) with sequenced genomes in the National Center for Biotechnology Information’s (NCBI) databases, supplemented with data on host habitat and energy source collected primarily from the Joint Genome Institute’s Genomes Online Database (JGI-GOLD). Extrachromosomal viruses represent ongoing acute or persistent infections in the microbial culture that was sequenced. Thus, increases in the axes labeled “… extrachromosomal viruses infecting” represent increasing viral coinfection in the host culture (not necessarily in single cells within that culture). The number of extrachromosomal viruses infecting a host was the response variable in a negative binomial regression that tested the effect of five variables (the number of prophage sequences, ssDNA virus presence, host taxon, host energy source, and habitat ecosystem) on variation in the response variable. Plots A-C depict all variables retained after stepwise model selection using Akaike’s Information Criterion (AIC). In Panel A, each point represents a microbial genus with its corresponding mean number of extrachromosomal virus infections (only genera with >1 host sampled and nonzero means are included). The colors of each point correspond to each genus. In Panels B and C, each point represents a unique host with a corresponding number of extrachromosomal virus infections. Panel B groups hosts according to their habitat ecosystem, as defined by JGI-GOLD database schema; the point colors correspond to each of these three ecosystem categories. Panel C groups hosts according to the number of prophage sequences detected in host sequences; the color shades of points become lighter in hosts with more detected prophages. Panels B and C have data points are offset by a random and small amount (jittered) to enhance visibility and reduce overplotting. The regression table is presented in Panel D, and only includes variables with p < 0.05.

### Single-cell coinfection is influenced by bacterial ecology, virus-virus interactions, and sequences associated with the CRISPR-Cas defense mechanism

To test the viral and host factors that affected viral coinfection of single cells, I constructed a Poisson regression on the single-cell coinfection data set, which characterized 143 viral infections in SUP-05 marine bacteria (sulfur-oxidizing Gammaproteobacteria) isolated as single cells (n=127) directly from their habitat (Roux *et al.,* 2014). Single cell coinfection was measured as the number of actively infecting viruses detected by sequence data in each cell. Stepwise model selection with AIC led to a model that included three factors explaining the number of actively infecting viruses in single cells: number of prophages, number of CRISPR spacers, and ocean depth at which cells were collected (Figure 3). The model estimated each additional prophage would result in ~9% of the amount of active infections (i.e. a 91% reduction), while each additional CRISPR spacer would result in a ~54% reduction. Depth had a positive influence on coinfection: every 50 meter increase in depth was predicted to result in ~80% more active infections.

**Figure 3.**
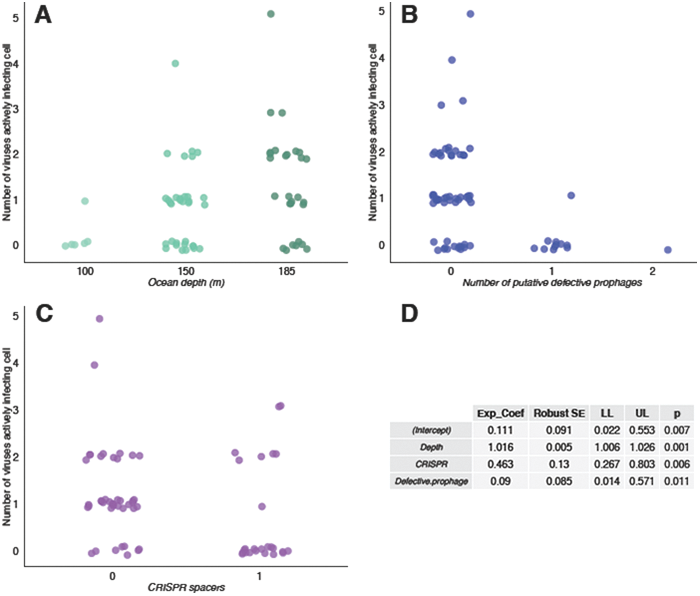
Host ecology, number of prophage sequences, and CRISPR spacers predict variation in single-cell coinfection. The data set is composed of viral infections (n=143) detected using genome sequence data in a set of SUP-05 (sulfur-oxidizing Gammaproteobacteria) marine bacteria isolated as single cells (n=127) from the Saanich Inlet in British Columbia, Canada. In Panels A-C each point represents a single-cell amplified genome (SAG). The y-axis depicts the number of viruses involved in active infections (active at the time of isolation, e.g. ongoing lytic infections) in each cell. Thus, increases in the y-axis represent increasing active viral coinfection of single cells. The number of viruses actively infecting each cell was the response variable in a Poisson regression that tested the effect of four variables (small subunit rRNA phylogenetic cluster, ocean depth, number of prophage sequences, and number of CRISPR spacers) on variation in the response variable. Panels A-C depict all variables retained after stepwise model selection using Akaike’s Information Criterion (AIC) and group cells according to the ocean depth at which the cells were isolated (A), the number of prophage sequences detected (B; conservatively termed putative defective prophages here, as in the original published dataset, because sequences were too small to be confidently assigned as complete prophages), and the number of CRISPR spacers detected (C); point colors correspond to the categories of each variable in each plot. Each point is offset by a random and small amount (jittered) to enhance visibility and reduce overplotting. The regression table is presented in panel D.

The phylogenetic cluster of the SUP-05 bacterial cells (based on small subunit rRNA amplicon sequencing) was not a statistically significant predictor selected using AIC model selection. Details of the full, reduced, and alternative models are provided in the Supplementary Materials and all associated code and data is deposited in FigShare repository.

### Prophages limit culture and single-cell coinfection

Based on the results of the regression analyses showing that prophage sequences limited coinfection, and aiming to test the phenomenon of superinfection exclusion in a large data set, I conducted a more detailed analysis of the influence of prophages on coinfection in microbial cultures. I also conducted a similar analysis using the single cell data set, which is relatively smaller (n=127 host cells), but provides better, single-cell level resolution.

First, because virus-virus interactions can occur in cultures of cells, I tested whether prophage-infected host cultures reduced the probability of other viral infections in the entire culture coinfection data set. A majority of prophage-infected host cultures were coinfections (56.48% of n = 3,134), a modest but statistically distinguishable increase over a 0.5 null (Binomial Exact: p = 4.344e^-13^, Figure 4A). Of these coinfected host cultures (n=1,770), cultures with more than one prophage sequence (32.54%) were two times less frequent than those with both prophage sequences and extrachromosomal viruses (Binomial Exact: p < 2.2e^-16^, Figure 4B). Therefore, integrated prophages appear to reduce the chance of the culture being infected with additional prophage sequences, but not additional extrachromosomal viruses. Accordingly, host cultures co-infected exclusively by extrachromosomal viruses (n=675) were infected by 3.25 ± 1.34 viruses, compared to 2.54 ± 1.02 prophage sequences (n=575); these quantities showed a statistically significant difference (Wilcoxon Rank Sum: W = 125398.5, p < 2.2e^-16^, Figure 4C).

**Figure 4.**
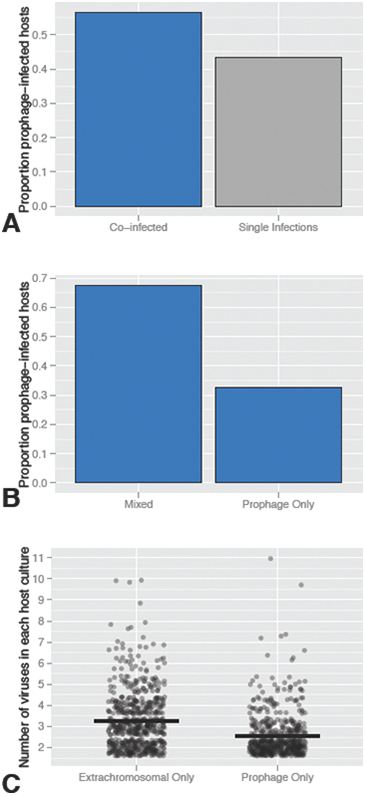
Host cultures infected with prophages limit coinfection by other prophages, but not extrachromosomal viruses. The initial data set was composed of viral infections detected using sequence data in sequenced genomes of microbial hosts available in National Center for Biotechnology Information (NCBI) databases. A slight, but statistically significant, majority of host cultures with prophage sequences (n = 3,134) were coinfected, i.e. had more than one extrachromosomal virus or prophage detected (Panel A, blue bar). Of these, host cultures containing multiple prophages were less frequent than those containing a mix of prophages and extrachromosomal (e.g. acute or persistent) infections (Panel B). On average (black horizontal bars), extrachromosomal-only coinfections involved more viruses than prophage-only coinfections (Panel C) In Panel C each point represents a bacterial or archaeal host and is offset by a random and small amount to enhance visibility.

Second, to test whether prophage sequences could affect coinfection of single cells in a natural environment, I examined a single cell amplified genomics data set of SUP05 marine bacteria. Cells with prophage sequences (conservatively termed putative defective prophages in the original published dataset, as in Figure 3B, because sequences were too small to be confidently assigned as complete prophages; S. Roux pers. comm) were less likely to have current infections: 9.09% of cells with prophage sequences had active viral infections, compared to 74.55% of cells that had no prophage sequences (X-squared = 14.2607, df = 1, p-value = 0.00015). Bacteria with prophage sequences were undergoing active infection by an average of 0.09 ± 0.30 phages, whereas bacteria without prophages were infected by 1.22 ± 1.05 phages (Figure 3B). No host with prophages had current coinfections (i.e., > 1 active virus infection).

### Non-random coinfection of host cultures by ssDNA and dsDNA viruses suggests mechanisms enhancing coinfection

The presence of ssDNA viruses in host cultures was not a statistically significant predictor of extrachromosomal infections in the regression analysis of the culture coinfection data set(based on Roux *et al.,* 2015) that is composed of sequence-based viral detections in all bacterial and archaeal NCBI genome sequences. However, to test the hypothesis that ssDNA-dsDNA viral infections exhibit non-random coinfection patterns (as found by Roux *et al.,* 2014 for single SUP05 bacterial cells), I conducted a more focused analysis on the culture coinfection data set. Coinfected host cultures containing ssDNA viruses (n = 331), were more likely to have dsDNA or unclassified viruses (70.69%), than multiple ssDNA infections (exact binomial: p= 3.314e^-14^). These coinfections were >2 times more likely to involve at least one dsDNA virus than none (exact binomial: p = 2.559e^-11^, Figure 5A). To examine whether differential detectability of ssDNA viruses could affect this result and place it in context of all the infections recorded in the data set, I conducted further tests. Single-stranded DNA viruses represented 6.82% of the viruses detected in host cultures in the overall data set (n=12,490). Coinfected host cultures that contained at least one ssDNA virus represented 6.19% of hosts, in line with, and not statistically different from the aforementioned overall detectability of ssDNA viruses (X^2^ = 3.22, df = 1, p-value = 0.07254). The most numerous coinfections in the culture coinfection data set were dsDNA-only coinfections (n=1,898). The proportion of ssDNA-mixed/total-ssDNA coinfections (0.71) was higher than the proportion of dsDNA-mixed/dsDNA-total (0.19) coinfections (Figure 5B; X^2^ = 398.55, df = 1, p < 2.2e^-16^). Of all the ssDNA viruses detected (n=852) in the data set, a majority belonged to the *Inoviridae* family (90.96%) with the remainder in the *Microviridae* (0.70%) or their family was unknown (8.33%). The proportion of hosts infected with *Inoviridae* was highest in dsDNA-ssDNA coinfections (0.96), higher than hosts with ssDNA-only coinfections (0.90) or than hosts with ssDNA single infections (0.86); the difference in these three proportions was statistically different from zero (X^2^ = 10.98, df = 2, p = 0.00413).

**Figure 5.**
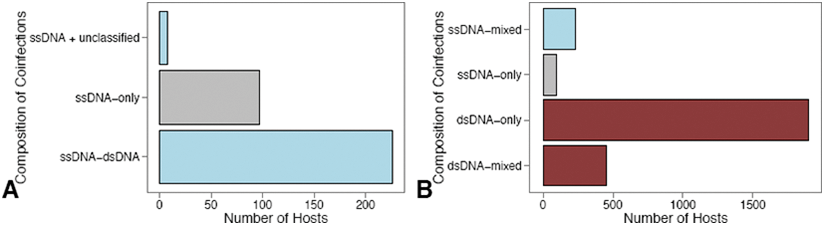
Culture coinfections between single-stranded DNA (ssDNA) and double stranded DNA (dsDNA) viruses are more common than expected by chance. The initial data set was composed of viral infections detected using sequence data in sequenced genomes of microbial hosts available in National Center for Biotechnology Information (NCBI) databases. Panel A: Composition of coinfections in all host cultures coinfected with at least one ssDNA virus (n=331) at the time of sequencing. Blue bars are mixed coinfections composed of at least one ssDNA virus and at least one dsDNA or unclassified virus. The grey bar depicts coinfections exclusively composed of ssDNA viruses (ssDNA-only). Host culture coinfections involving ssDNA viruses were more likely to occur with dsDNA or unclassified viruses, than with multiple ssDNA viruses (exact binomial: p= 3.314e^-14^). Panel B: The majority of host culture coinfections in the data set exclusively involved dsDNA viruses (n= 1,898; dsDNA-only red bar). There were more mixed coinfections involving ssDNA viruses than ssDNA-only coinfections (compare blue and grey bar), while dsDNA-only coinfections were more prevalent than mixed coinfections involving dsDNA viruses (compare red bars). Accordingly, the proportion of ssDNA-mixed/total-ssDNA coinfections was higher than the proportion of dsDNA-mixed/dsDNA-total coinfections (proportion test: p < 2.2e^-16^).

### CRISPR spacers limit coinfection at single cell level without spacer matches

The regression analysis of the single-cell data set (n=127 cells of SUP-05 marine bacterial cells) revealed that the presence of CRISPR spacers had a significant overall effect on coinfection of single cells; therefore I examined the effects of CRISPR spacers more closely. Cells with CRISPR spacers were less likely to have active viral infections than those without spacers (X-squared = 14.03, df = 1, p-value = 0.00018). Cells with CRISPR spacers were less likely to have active viral infections, with 32.00% of cells with CRISPRs having active viral infections, compared to 80.95% percent of bacteria without CRISPR spacers. Bacterial cells with CRISPR spacers had 0.68 ± 1.07 current phage infections compared to 1.21 ± 1.00 for those without spacers (Figure 3C). Bacterial cells with CRISPR spacers could have active infections and coinfections with up to 3 phages. None of the CRISPR spacers matched any of the actively infecting viruses (Roux *et al.,* 2014), using an exact match search (S. Roux, pers. comm.).

## Discussion

### Summary of findings

The compilation of multiple large-scale data sets of virus-host interactions (>6,000 hosts, >13,000 viruses) allowed a systematic test of the ecological and biological factors that influence viral coinfection rates in microbes across a broad range of taxa and environments. I found evidence for the importance and consistency of host ecology and virus-virus interactions in shaping potential (cross-infectivity), culture, and single-cell coinfection. In contrast, host taxonomy was a less consistent predictor of viral coinfection, being a weak predictor of cross-infectivity and single cell coinfection, but representing the strongest predictor of host culture coinfections. In the most comprehensive test of the phenomenon of superinfection immunity conferred by prophages (n = 3,134 hosts), I found that prophage sequences were predictive of limited coinfection of cultures by other prophages, but less strongly predictive of coinfection by extrachromosomal viruses. Furthermore, in a smaller sample of single cells of SUP-05 marine bacteria (n=127 cells), prophage sequences completely excluded coinfection (by prophages or extrachromosomal viruses). In contrast, I found evidence of *increased* culture coinfection by ssDNA and dsDNA phages, suggesting mechanisms that may enhance coinfection. At a fine-scale, single-cell data revealed that CRISPR spacers limit coinfection of single cells in a natural environment, despite the absence of exact spacer matches in the infecting viruses, suggesting a potential role for bacterial defense mechanisms. In light of the increasing awareness of the widespread occurrence of viral coinfection, this study provides the foundation for future work on the frequency, mechanisms, and dynamics of viral coinfection and its ecological and evolutionary consequences.

### Host correlates of coinfection

Host ecology stood out as an important predictor of coinfection across all three datasets: cross-infectivity, culture coinfection, and single cell coinfection. The specific ecological variables differed in the data sets (bacterial association, isolation habitat, and ocean depth), but ecological factors were retained as statistically significant predictors in all three models. Moreover, when ecological variables were standardized between the cross-infectivity and culture coinfection data sets, the differences in cross-infectivity and coinfection among ecosystem categories were remarkably similar (Figure S5, Table S2). This result implies that the ecological factors that predict whether a host *can* be coinfected are the same that predict whether a host *will* be coinfected. These results lend further support to the consistency of host ecology as a predictor of coinfection. On the other hand, host taxonomy was a less consistent predictor. It was weak or absent in the cross-infectivity and single-cell coinfection models, respectively, yet it was the strongest predictor of culture coinfection. This difference could be because the hosts in the cross-infectivity and single-cell data sets varied predominantly at the strain level (Flores *et al.,* 2011; Roux *et al.,* 2014), whereas infection patterns are less variable at the Genus level and higher taxonomic ranks (Flores *et al.,* 2013; Roux *et al.,* 2015) as seen with the culture coinfection model. Collectively, these findings suggest that the diverse and complex patterns of cross-infectivity (Holmfeldt *et al.,* 2007) and coinfection observed at the level of bacterial strains may be best explained by local ecological factors, while at higher taxonomic ranks the phylogenetic origin of hosts increases in importance (Flores *et al.,* 2011; 2013; Roux *et al.,* 2015). Particular bacterial lineages can exhibit dramatic differences in cross-infectivity (Koskella and Meaden, 2013; Liu *et al.,* 2015) and, as this study shows, coinfection. Thus, further studies with ecological data and multi-scale phylogenetic information will be necessary to test the relative influence of bacterial phylogeny on coinfection.

Sequences associated with the CRISPR-Cas bacterial defense mechanism were another important factor influencing coinfection patterns, but were only tested in the comparatively smaller single-cell data set (n=127 cells of SUP05 marine bacteria). The presence of CRISPR spacers reduced the extent of active viral infections, even though these spacers had no exact matches (S. Roux, pers. comm.) to any of the infecting viruses identified (Roux *et al.,* 2014). These results provide some of the first evidence from a natural environment that CRISPR’s protective effects could extend beyond viruses with exact matches to the particular spacers within the cell (Fineran *et al.,* 2014; Semenova *et al.,* 2011). Although very specific sequence matching is thought to be required for CRISPR-Cas-based immunity (Barrangou *et al.,* 2007; Brouns *et al.,* 2008; Mojica *et al.,* 2005), the system can tolerate mismatches in protospacers (within and outside of the seed region: Semenova *et al.,* 2011; Fineran *et al.,* 2014), enabling protection (interference) against related phages by a mechanism of enhanced spacer acquisition termed priming (Fineran *et al.,* 2014). The seemingly broader protective effect of CRISPR-Cas beyond specific sequences may help explain continuing effectiveness of CRISPR-Cas (Fineran *et al.,* 2014) in the face of rapid viral coevolution for escape (Heidelberg *et al.,* 2009; Tyson and Banfield, 2008; Andersson and Banfield, 2008). However, caution is warranted as CRISPR spacers sequences alone cannot indicate whether CRISPR-Cas is active. Some bacterial taxa have inactive CRISPR-Cas systems, notably *Escherichia coli K-12* (Westra *et al.,* 2010), and phages also possess anti-CRISPR mechanisms (Bondy-Denomy *et al.,* 2015; Seed *et al.,* 2013) that can counter bacterial defenses. In this case the unculturability of SUP05 bacteria preclude laboratory confirmation of CRISP-Cas activity, but given the strong statistical association of CRISPR space sequences with a reduction in active viral infections (after controlling for other factors), it is most parsimonious to tentatively conclude that the CRISPR-Cas mechanism is the likeliest culprit behind these reductions. To elucidate and confirm the roles of bacterial defense systems in shaping coinfection, more data on CRISPR-Cas and other viral infection defense mechanisms will be required across different taxa and environments.

This study revealed the strong predictive power of several host factors in explaining viral coinfection, yet there was still substantial unexplained variation in the regression models (see Results). Thus, the host factors tested herein should be regarded as starting points for future experimental examinations. For instance, host energy source was not a statistically significant predictor in the cross-infectivity and culture coinfection models, perhaps due to the much smaller sample sizes of autotrophic hosts. However, both data sets yield similar summary statistics, with heterotrophic hosts having 59% higher cross-infectivity and 77% higher culture coinfection (Figure S6). Moreover, other factors not examined, such as geography could plausibly affect coinfection. The geographic origin of strains can affect infection specificity such that bacteria isolated from one location are likely to be infected by more phage isolated from the same location, as observed with marine microbes (Flores *et al.,* 2013). This pattern could be due to the influence of local adaptation of phages to their hosts (Koskella *et al.,* 2011) and represents an interesting avenue for further research.

### The role of virus-virus interactions in coinfection

The results of this study suggest that virus-virus interactions play a role in limiting *and* enhancing coinfection. First, host cultures and single cells with prophage sequences showed limited coinfection. In what is effectively the largest test of viral superinfection exclusion (n = 3,134 hosts), host cultures with prophage sequences exhibited limited by other prophages, but not as much by extrachromosomal viruses. Although this focused analysis showed that prophage sequences limited extrachromosomal infection less strongly than additional prophage infections, the regression analysis suggested that they did have an effect: a statistically significant ~21% reduction in extrachromosomal infections for each additional prophage detected. As these were culture coinfections and not necessarily single-cell coinfections, these results are consistent with a single-cell study of *Salmonella* cultures showing that lysogens can activate cell subpopulations to be transiently immune from viral infection (Cenens *et al.,* 2015). Prophage sequences had a more dramatic impact at the single-cell level in SUP05 marine bacteria in a natural environment, severely limiting active viral infection and completely excluding coinfection. The results on culture-level and single-cell coinfection come from very different data sets, which should be examined carefully before drawing general patterns: The culture coinfection data set is composed of an analysis of all publicly available bacterial and archaeal genome sequences in NCBI databases that show evidence of a viral infection. These sequences show a bias towards particular taxonomic groups (e.g. *Proteobacteria,* model study species) and those that are easy to grow in pure culture. The single cell data set is limited to 127 cells of just one host type (SUP05 marine bacteria) isolated in a particular environment, as opposed to the 5,492 hosts in the culture coinfection data set. This limitation prohibits taxonomic generalizations about the effects on prophage sequences on single cells, but extends laboratory findings to a natural environment. Additionally, both of these data sets use sequences to detect the presence of prophages, which means the activity of these prophage sequences cannot be confirmed, as with any study using sequence data alone. Along these lines, in the original single cell coinfection data set (Roux *et al.,* 2014), the prophage sequences were conservatively termed ‘putative defective prophages’ (as in Figure 3 here), because sequences were too small to be confidently assigned as complete prophage sequences (S. Roux, pers. comm.). If prophages in either data set are not active, this could mean that bacterial domestication of phage functions (Bobay *et al.,* 2014; Asadulghani *et al.,* 2009), rather than phage-phage interactions in a strict sense, would explain protection from infection conferred by these prophage sequences in host cultures and single cells. In view of these current limitations, a wider taxonomic and ecological range of culture and single-cell sequence data, together with laboratory studies when possible, should confirm and elucidate the role of prophage sequences in affecting coinfection dynamics. Interactions in coinfection between temperate bacteriophages can affect viral fitness (Dulbecco, 1952; Refardt, 2011), suggesting latent infections are a profitable avenue for future research on virus-virus interactions.

Second, another virus-virus interaction examined in this study appeared to increase the chance of coinfection. While prophage sequences strongly limited coinfection in single cells, Roux et al.’s (2014) original analysis of this same data set found strong evidence of enhanced coinfection (i.e. higher than expected by random chance) between dsDNA and ssDNA *Microviridae* phages in bacteria from the SUP05_03 cluster. I conducted a larger test of this hypothesis using a different, diverse set of 331 bacterial and archaeal hosts infected by ssDNA viruses included in the culture coinfection data set (Roux *et al.,* 2015). This analysis provides evidence that ssDNA-dsDNA culture coinfections occur more frequently than would be expected by chance, extending the taxonomic applicability of this result from one SUP05 bacterial lineage to hundreds of taxa.

Thus, enhanced coinfection, perhaps due to the long, persistent infection cycles of some ssDNA viruses, particularly the *Inoviridae* (e.g. Innoviridae: Rakonjac *et al.,* 2011), might be a major factor explaining findings of phages with chimeric genomes composed of different types of nucleic acids (Roux *et al.,* 2013; Diemer and Stedman, 2012). Filamentous or rod-shaped ssDNA phages (family Inoviridae; Székely and Breitbart, 2016) often conduct multiple replications without killing their bacterial hosts (Rakonjac *et al.,* 2011; Székely and Breitbart, 2016; Mai-Prochnow *et al.,* 2015), allowing more time for other viruses to coinfect the cell and providing a potential mechanistic basis for the enhanced dsDNA-ssDNA coinfections. In line with this prediction, over 96% of dsDNA-ssDNA host culture coinfections contained at least one virus from the *Inoviridae*, a greater (and statistically distinguishable) percentage than contained in ssDNA-only coinfections or ssDNA single infections. Notwithstanding, caution is warranted due to known biases against the detectability of ssDNA compared to dsDNA viruses in high-throughput sequencing library preparation (Roux *et al.,* 2016). In this study, this concern can be addressed by noting that: not all the sequencing data in the culture coinfection data set was generated by high-throughput sequencing, the prevalence of host coinfections containing ssDNA viruses was virtually identical (within 0.63%) to the detectability of ssDNA viruses in the overall data set, and the prevalence of ssDNA viruses in this data set (~6%) is in line with validated, unbiased estimates of ssDNA virus prevalence from natural (marine) habitats (Roux *et al.,* 2016). However, given that the ssDNA virus sample is comparatively smaller than the dsDNA sample, future studies should focus on examining ssDNA-dsDNA coinfection prevalence and mechanisms.

Collectively, these results highlight the importance of virus-virus interactions as part of the suite of evolved viral strategies to mediate frequent interactions with other viruses, from limiting to promoting coinfection, depending on the evolutionary and ecological context (Turner and Duffy, 2009).

### Implications and applications

In sum, the results of this study suggest microbial host ecology and virus-virus interactions are important drivers of the widespread phenomenon of viral coinfection. An important implication is that virus-virus interactions will constitute an important selective pressure on viral evolution. The importance of virus-virus interactions may have been underappreciated because of an overestimation of the importance of superinfection exclusion (Dulbecco, 1952). Paradoxically, superinfection avoidance may actually highlight the selective force of virus-virus interactions. In an evolutionary sense, this viral trait exists precisely because the potential for coinfection is high. If this is correct, variability in potential and realized coinfection, as found in this study, suggests that the manifestation of superinfection exclusion will vary across viral groups according to their ecological context. Accordingly, there are specific viral mechanisms that promote coinfection, as found in this study with ssDNA/dsDNA coinfections and in other studies (Dang *et al.,* 2004; Cicin-Sain *et al.,* 2005; Turner *et al.,* 1999; Altan-Bonnet and Chen, 2015; Erez *et al.,* 2017; Aguilera *et al.,* 2017). I found substantial variation in cross-infectivity, culture coinfection, and single-cell coinfection and, in the analyses herein, ecology was always a statistically significant and strong predictor of coinfection, suggesting that the selective pressure for coinfection is going to vary across local ecologies. This is in agreement with observations of variation in viral genetic exchange (which requires coinfection) rates across different geographic localities in a variety of viruses (Díaz-Muñoz *et al.,* 2013; Trifonov *et al.,* 2009; Held and Whitaker, 2009).

These results have clear implications, not only for the study of viral ecology in general, but for practical biomedical and agricultural applications of phages and bacteria/archaea. Phage therapy is often predicated on the high host specificity of phages, but intentional coinfection could be an important part of the arsenal as part of combined or cocktail phage therapy. This study also suggests that viral coinfection in the microbiome should be examined, as part of the influence of the virome on the larger microbiome (Pride *et al.,* 2012; Minot *et al.,* 2011; Reyes *et al.,* 2010). Finally, if these results apply in eukaryotic viruses and their hosts, variation in viral coinfection rates should be considered in the context of treating and preventing infections, as coinfection likely represents the default condition of human hosts (Wylie *et al.,* 2014). Coinfection and virus-virus interactions have been implicated in changing disease outcomes for hosts (Vignuzzi *et al.,* 2006), altering epidemiology of viral diseases (Nelson *et al.,* 2008), and impacting antimicrobial therapies (Birger *et al.,* 2015). In sum, the results of this study suggest that the ecological context, mechanisms, and evolutionary consequences of virus-virus interactions should be considered as an important subfield in the study of viruses.

## Acknowledgements

Simon Roux kindly provided extensive assistance with previously published data sets. T.B.K. Reddy kindly provided database records from Joint Genome Instituteߣs Genomes Online Database. I am indebted to Joshua Weitz, Britt Koskella, and Jay Lennon for providing helpful critiques and advice on an earlier version of this manuscript. A Faculty Fellowship to SLDM from New York University supported this work.

## Data availability

A list and description of data sources are included in Supplementary Table 1, and the raw data and code used in this paper are deposited in the FigShare data repository (FigShare doi:10.6084/m9.figshare.2072929).

